# Single-molecule interaction microscopy reveals antibody binding kinetics

**DOI:** 10.1101/2020.09.21.306605

**Authors:** Thilini Perera, Hirushi Gunasekara, Ying S. Hu

**Affiliations:** Department of Chemistry, College of Liberal Arts and Sciences, University of Illinois at Chicago, Chicago, IL, 60607-7061, USA

## Abstract

Single-molecule imaging has provided new insights on weak transient biomolecular interactions with micromolar to millimolar affinity. However, the limited duration of observation has hindered the study of strong and reversible interactions with sub-nanomolar affinity. We report single-molecule interaction microscopy (SMIM), which combines point accumulation for imaging in nanoscale topography (PAINT) with extended imaging durations that enables the study of antibody binding kinetics in the cellular environment. SMIM revealed heterogeneous binding kinetics and the effect of concentration and antibody valency on the association and dissociation rates on antibody-antigen interactions in their cellular environments. We thereby demonstrate SMIM as a versatile single-molecule technique for studying strong, transient biomolecular interactions.

---

Protein-protein interactions comprise the fundamental mechanisms for a number of biological processes ranging from enzyme catalysis to regulation of immune responses. These interactions are transient, having a wide range of affinities, and can be generally classified as strong or weak interactions ^1^. The transient nature of these interactions facilitated by reversible binding enables biological organisms to rapidly respond to highly dynamic cellular processes ^2,3^. Among different classes of transient interactions in biological systems, the equilibrium dissociation constant, K_D_, ranges from 10^−7^ to 10^−3^ M for weak enzyme-substrate interactions to 10^−12^ to 10^−8^ M for strong antibody-antigen interactions ^4,5^. The K_D_ values obtained by ensemble-averaged techniques, such as surface plasmon resonance ^6^, biolayer interferometry ^7^ and microscale thermophoresis ^8^, however, do not characterize the dynamic process of these interactions in the native environment of the cell. In some cases, relatively large amounts of the reagents, such as antibodies and binding partners, are costly or difficult to produce.

Recent advancements in single-molecule imaging have improved our understanding of weak transient interactions, including the binding of transcriptional factors to DNAs ^9,10^, ribosomes to mRNAs ^11,12^, and signaling molecules to membrane proteins and lipids ^13–15^. Despite these new developments, imaging tools to study strong-transient interactions at the single-molecule level remain scarce. For instance, the strong-transient antibody-antigen interactions occur at a time scale that exceeds the observation period of single molecules, which is typically limited to a few seconds due to photobleaching. Several sample treatment strategies to prolong the duration of single-molecule imaging have been reported, including deoxygenation ^16,17^ and the use of a reducing-plus-oxidizing system ^18–20^. However, the utilization of these methods is limited due to their toxicity to living cells. Here, we introduce single-molecule interaction microscopy (SMIM) as a potential solution. SMIM merges the point accumulation for imaging in nanoscale topography (PAINT) technique ^21–23^ with extended imaging duration, thereby enabling the investigation of antibody binding kinetics in the cellular environment without sample manipulation.

SMIM achieves extended imaging durations by inserting non-illuminating intervals (NII) between consecutive image frames **(Fig. 1a)**. The duration of NII was progressively increased until distinct binding kinetics were observed. By this method, the fluorophore lifetime could be stretched over an arbitrary period ^24^. **Fig. 1b** demonstrates a single-molecule intensity trajectory of 287 s recorded using a 5 s NII from immobilized Superclonal^™^ goat anti-rabbit Alexa Fluor^™^ 647 (AF 647) antibodies on a clean cover glass. Under continuous illumination, single-molecule intensity trajectories lasted for an average of only 2.7 s until photobleaching (Methods). The camera exposure time was kept at 50 ms to achieve a sufficient signal-to-background ratio. The stepwise photobleaching of individual antibody molecules was consistent with the ensemble measurements (**Supplementary Fig. 1a**). **Fig. 1c** shows that the extended imaging duration correlated with the length of the NII. The insertion of 0.5 s and 5 s NIIs extended the average imaging duration to 26 s and 234 s (**Fig. 1c**), or a fold increase of 9.63 and 86.67, respectively. SMIM employs a customized MATLAB code (Methods) from a single-molecule tracking code for visualization ^25^. The program displays single-molecule events as color-coded circles with different radii corresponding to their dwell times (**Fig. 1d**).

**Figure 1.**
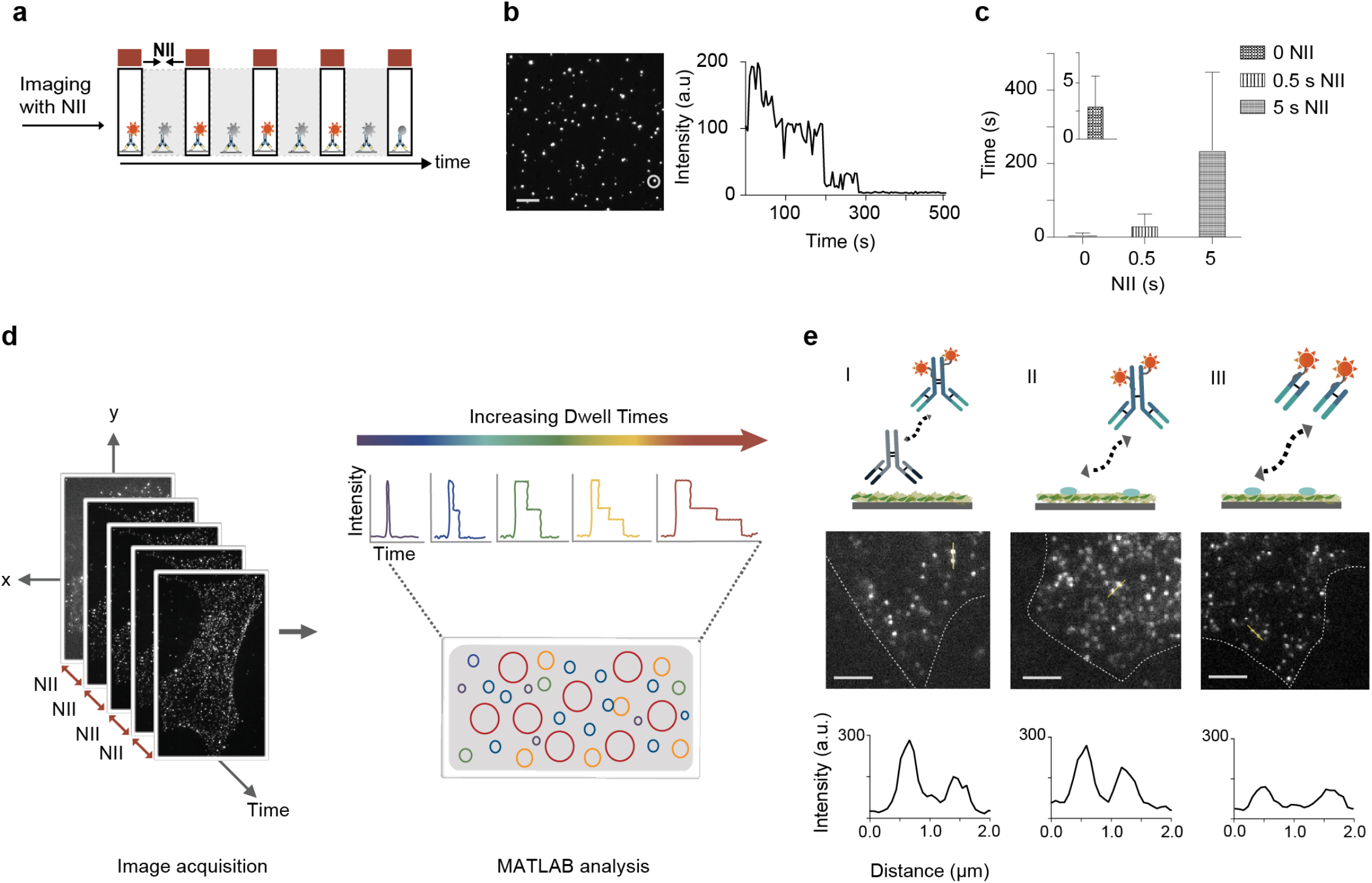
The principle of SMIM as a versatile single-molecule technique to study strong transient molecular interactions with extended imaging durations. **a** Schematic representation of the imaging scheme with non-illuminating intervals (NII). **b**, Single-molecule images from a cover glass and a corresponding extended single-molecule fluorescence trajectory. **c**, Extended imaging durations with different NIIs. **d**, Schematic representation of SMIM workflow to study antibody binding kinetics. Following sample preparation, the image sequence was acquired with the chosen NII and analyzed with a custom-coded single-particle tracking MATLAB program. **e**, Selected antibody-interaction conditions studied, **I**, secondary antibody interaction (Superclonal^™^ goat-anti-rabbit IgG-AF647 with rabbit anti-tubulin primary antibody), **II**, primary antibody (12CA5 IgG-AF647 with HA) and **III**, Fab (Fab-AF647 with HA) interactions with the subcellular target. (Row 1) and the corresponding single-molecule images and signal-to-noise ratios (Row 2 and Row 3). The image sequence in **b** and **d** was acquired with a 5 s NII. Scale bar: 5 µm.

The strong transient nature of antibody-antigen interactions depends on a multitude of factors, among which the affinity, avidity, and kinetics of the interactions play a major role ^26^. We used SMIM to investigate the antibody binding kinetics in their cellular environment (**Fig. 1e, top row**). In the first case, we studied the concentration dependent binding kinetics of a commercial Superclonal^™^ secondary antibody to the anti-tubulin primary antibody. In the second and third case, we investigated the direct interactions of bivalent and monovalent forms of anti-HA antibody to the HA-tag to study the effect of antibody valency as a contributing factor to the overall strength of the antibody-antigen complex dependent on the structural arrangement of the interacting partners ^27–29^. To this end, we generated cell lines expressing three hemagglutinin (HA) tags on the N terminus of the α-tubulin and prepared fluorescently labeled anti-HA antibodies (12CA5 IgG) and monovalent 12CA5 antibody fragments (Fab) for SMIM. With the same exposure time as in **Fig 1b,c**, a negligible background from freely diffusing dye labeled antibodies were observed for each case studied (**Fig. 1e**). By combining a suitable NII with a simple PAINT approach on antibodies, SMIM enables visualization of single-molecule antibody-antigen interactions beyond conventional dwell times.

## Results

### SMIM revealed heterogeneous binding kinetics for antibody-antigen interactions

We first investigated the binding behavior of Superclonal^™^ IgG antibodies in the presence of immobilized primary antibodies against the β-tubulin in U2OS cells. Superclonal^™^ antibodies are recombinant polyclonal secondary antibodies engineered to achieve multi-epitope coverage. **Fig. 2a,b** illustrate heterogeneous binding dynamics of the antibody at 0.1 nM. The dwell times of binding events were calculated by the number of frames with the corresponding NII. We confirmed no significant correlation between the dwell time and brightness of antibody molecules by performing Pearson’s analysis (Methods, Pearson’s *r* of 0.1436 and p <0.0001, **Supplementary Fig. 2a)**. At 0.1 nM concentration, SMIM revealed a higher abundance of stable interactions (greater than 70 s, large, red circles) compared to transient interactions (less than 30 s, small, purple circles). Representative single-molecule intensity profiles corresponding to these transient (narrow peaks, upper panels) and stable (broader peaks, lower panels) interactions are shown in **Fig. 2c** and **Supplementary Fig. 3**. Additional data indicated that subsequent stable binding could occur in the same diffraction-limited site following initial transient interactions (**Supplementary Fig. 4**).

**Figure 2.**
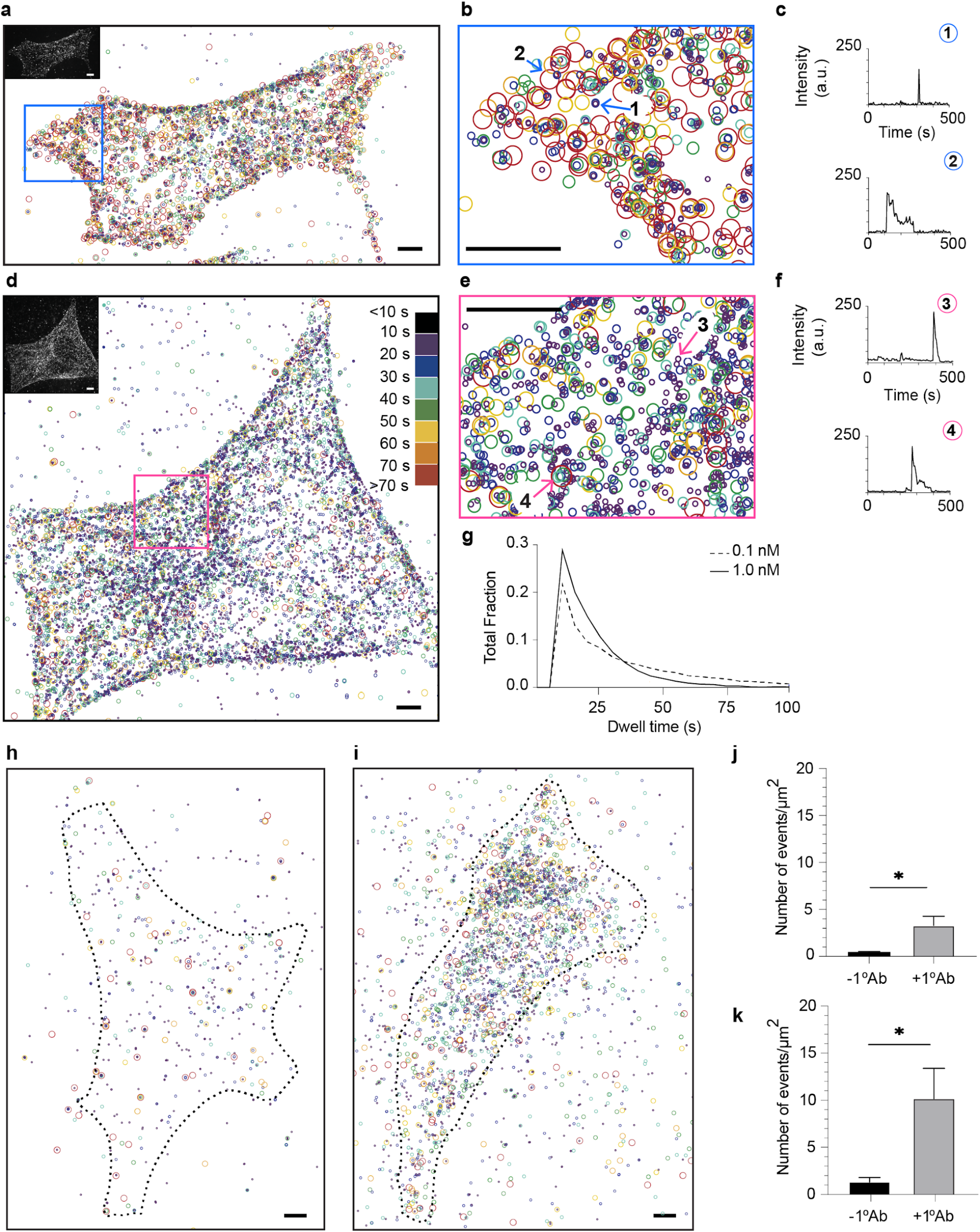
Visualizing heterogeneous, concentration-dependent acceleration of binding kinetics of secondary antibody interactions in a cellular environment with SMIM. SMIM interaction maps of Superclonal^™^ goat anti-rabbit IgG-Alexa Fluor^™^ 647 (AF647) at **a**, 0.1 nM, **d**, 1.0 nM in a fixed U2OS cell labeled with the corresponding anti-tubulin primary antibody. **b, e**, High-magnification views of the boxed regions in **a** and **d**, respectively. **c, f** representative single-molecule intensity plots of transient (1,3) and stable binding events (2,4) for the selected events in **b** and **e. g**, Statistical analysis of binding events in secondary antibody staining at 0.1 nM 1.0 nM. SMIM interaction map showing color-coded dwell times of Superclonal^™^ goat anti-rabbit IgG-Alexa Fluor^™^ 647 (AF647), **h**, 0.1 nm, **i**, 1.0 nM, in a fixed U2OS without the primary antibody labeling; Cell shape is contoured. Comparison of single-molecule events from Superclonal^™^ goat anti-rabbit IgG-AF647 antibodies interacting with a fixed U2OS cell with (+1°Ab) and without (−1°Ab) the primary antibody labeling at **j**, 0.1 nM and **k**, 1.0 nM concentrations. The error bar represents the SD of the cell-to-cell variation. Statistical significance was evaluated using an unpaired Student’s t-test and *represents *p<*0.05 (*n*=5). Scale bars: 5 µm

### SMIM illustrates the concentration-dependent association/dissociation (on/off) rates of antibody kinetics

At a 10-fold higher concentration of 1 nM, SMIM revealed binding kinetics of the Superclonal^™^ IgG. The observed binding kinetics dramatically shifted to small, purple circles, signifying an increased presence of transient interactions (**Fig. 2d,e). Fig. 2f** illustrates representative single-molecule intensity profiles corresponding to transient (upper panels) and stable (lower panels) interactions. Statistical analysis demonstrated that the fraction of single-molecule events with dwell times less than 30 s increased at 1.0 nM antibody concentration (**Fig. 2g**). This observation suggests a competitive nature for antibody kinetics at higher concentrations that shifts the overall binding dynamics towards transient interactions. To confirm that our observations were due to the specific binding of the secondary antibody to the primary antibody, we performed SMIM in the absence of a primary antibody. SMIM on cells without the primary antibody exhibited a relatively small number of scattered and non-specific interactions (**Fig 2h, i**). **Fig 2j,k** compare the total binding event densities with and without primary antibody at 0.1 nM and 1.0 nM, respectively. We calculated the specific binding density by subtracting the non-specific interactions from total binding events. The resulting specific binding density increased three-fold at 1 nM over 0.1 nM. Despite the fact that SMIM registers the newly bound antibody molecules, which only accounts for the non-photobleached fraction of the total bound antibody molecules, the data are consistent with ensemble kinetics in which the on rate of the interaction is directly related to the antibody concentration.

Next, we applied SMIM to study the binding kinetics of the monoclonal 12CA5 antibody against the nine-amino-acid HA tag. We generated U2OS cell lines from single-cell clones (2E8 and 2F6) that express 3xHA-α-tubulin. Commercial 12CA5 antibodies were labeled with Alexa Fluor^™^ 647 (AF647) and used for SMIM experiments at 0.1 and 1.0 nM (**Supplementary Fig. 5a-c**). Pearson’s analysis confirmed no correlation between the measured dwell times and the initial brightness of the antibody molecule (Pearson’s r = 0.0627 and p <0.0001, **Supplementary Fig. 2b**). At both concentrations, we observed similar heterogenous and concentration-dependent binding kinetics for 12CA5 IgG-AF647 as we did with Superclonal^™^ IgG-AF647. It is intriguing that the microtubule structure started to emerge for both Superclonal^™^ IgG antibodies (**Fig. 2d**) and 12CA5 IgG-AF647 (**Fig. 3a**) from 100 frames of SMIM interaction maps at 1 nM while the majority of the binding events manifested as transient events. Hence, SMIM enables successful visualization of antibody interaction kinetics at different concentrations in the cellular environment.

**Figure 3.**
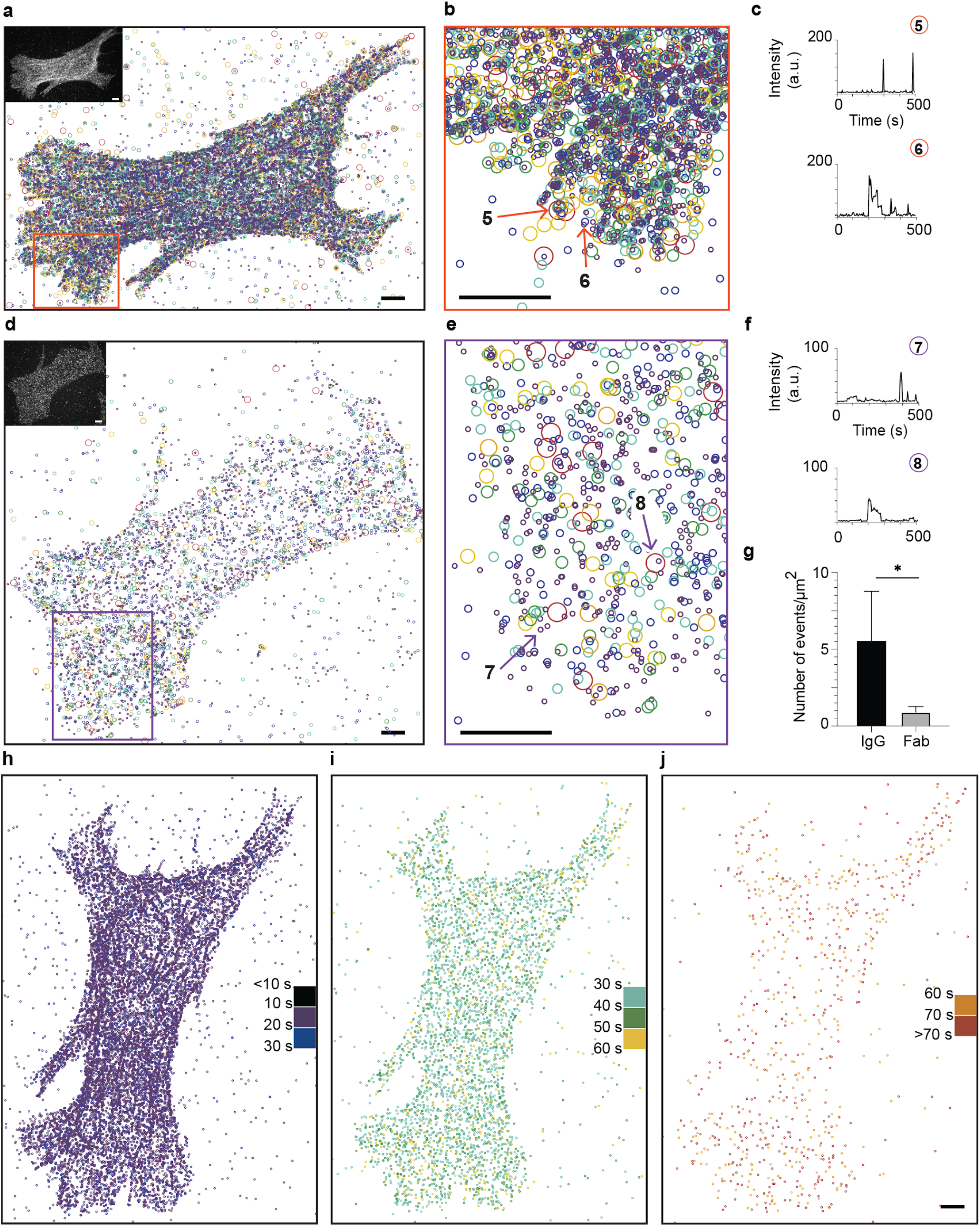
SMIM illustrates the effect of antibody valency on antibody-antigen interactions in a cellular environment with SMIM. SMIM interaction maps of primary staining with 12CA5 IgG-AF647 (**a**) and 12CA5 Fab-AF647 (**d)** each at 1.0 nM in a fixed mutant U2OS cell (2F6) expressing 3xHA-α-tubulin. **b, e**, High-magnification views of the boxed regions in **a** and **d** respectively. **c, f**, representative single-molecule intensity plots of transient (5,7) and stable binding events (6,8) for the selected events in **b** and **e. g**, Statistical analysis of specific binding event densities between 12CA5 IgG-AF647 and 12CA5 Fab-AF647 staining, each at 1.0 nM concentration. The error bar represents the SD of the cell-to-cell variation. Statistical significance was evaluated using an unpaired Student’s t-test and *represents *p<*0.05 (*n*=5). Dwell time breakdown of SMIM map of the panel **a**, (rotated 90° for presentation) illustrating interacting events with dwell times less than 30 seconds, **h**, between 30-60 seconds, **i**, and greater than 60, **j**. Scale bars: 5 µm

### SMIM illustrates the effect of antibody valency on interaction kinetics

To study the effect of antibody valency on the binding kinetics, we generated dye-labeled monovalent 12CA5 antibody fragments (12CA5 Fab-AF647) by enzymatic cleavage of 12CA5 IgG-AF647 using papain (**Methods, Supplementary Fig. 6**). The HA expression on the α-tubulin was confirmed by immunofluorescence staining using 12CA5 IgG-AF647 **(Supplementary Fig. 7**). **Fig. 3 a,b** and **d,e** show the SMIM interaction maps of the 12CA5 IgG-AF647 and 12CA5 Fab-AF647, respectively. We observe a distinct reduction in the binding density of monovalent 12CA5 Fab-AF647 compared to its divalent counterpart 12CA5 IgG-AF647 at 1 nM (**Fig. 3a** vs **d** and **Supplementary Fig**.**5a vs. d). Fig. 3c,f** show time trajectories of representative transient (upper panels) and stable (lower panels) binding events. The single-Fab intensity levels were approximately one-third of the IgG tracks, consistent with its reduced size compared to the full-length counterpart. Furthermore, the characteristic single-molecule intensity profiles remain consistent between IgG and Fab. (**Fig. 3c *vs*. 3f)**. SMIM maps of both 12CA5 IgG-AF647 and 12CA5 Fab-AF647 exhibit similar statistics of binding events (**Supplementary Fig. 5c and f**), implying similar binding dynamics between the IgG and the Fab. However, SMIM reveals a six-fold reduction in the binding density of 12CA5-Fab compared to 12CA5-IgG with no apparent microtubule structure at 1.0 nM concentration **(Fig. 3g, Supplementary Fig. 5a vs. d**). Since binding density is a direct indication of the on-rate, this suggests a six-fold reduction of the on-rate and thereby the overall affinity for 12CA5-Fab. This observation illustrates the significance of binding valency on interaction kinetics. We further experimented the effect of concentration on the dwell time distributions and on/off rates of 12CA5 Fab-AF647 (**Supplementary Fig 5d-f)**. The monovalent Fab at both 0.1 nM and 1.0 nM concentrations exhibited similar interaction kinetics. In addition, no difference in binding densities was found between 12CA5 IgG-AF647 and 12CA5 Fab-AF647 at 0.1 nM concentration (**Supplementary Fig. 5b and e**). This observation further reflects the reduction in affinity and association rates with antibody valency.

### SMIM reveals transient but specific interactions

Intuitively, specific interactions have longer dwell times (stable interactions), while non-specific or rather less-specific interactions tend to have shorter dwell times (transient interactions). However, SMIM reveals specific yet transient transitions. **Fig. 3 h-j** shows the breakdown of dwell-time distributions of 12CA5 IgG-AF647 of **Fig 3a** into three groups, less than 30 s, between 30-60 s and greater than 60 seconds. The marker size of the individual events was kept the same in all panels for clarity. The locations of specific interactions marked by the microtubule structure were mostly outlined by short dwell times (transient interactions) compared to events with longer dwell times **Fig. 3h** vs. **i,j**. This trend was also visible with Superclonal^™^-AF647 at 1.0 nM concentration. At concentrations sufficient to drive the binding kinetics close to equilibrium, antibody-antigen interactions are reversible and the dwell times take a turn from a long time regime to shorter times with the additional influence from the constant competition. SMIM data acquired over an extended period confirms constant and reversible binding kinetics of the 12CA5-IgG antibodies. **Supplementary Fig. 8** illustrates the binding kinetics over 100 min after complete photobleaching of a fluorescently labeled cell by 12CA-IgG AF647. The initial fluorescent recovery was registered by a relatively low density of stable binding events (**Supplementary Fig. 8 a,b**). The later frames were marked by high densities of more transient binding events and the kinetics remained constant after reaching the equilibrium (**Supplementary Fig. 8 c-l**). Thus, SMIM unravels subtle and complex antibody-antigen interaction dynamics that had not been observed by other imaging techniques.

## Discussion

We developed SMIM as a versatile single-molecule technique for the characterization of strong transient interactions between antibodies and immobilized binding partners. SMIM does not require special reagents or instrumentation and is useful for a wide range of studies, such as protein-protein interactions and small-molecule binding assays for drug discovery. SMIM circumvents the short bias in single-molecule tracking due to photobleaching. By progressively increasing the NII, an unbiased track length can be determined at the mild cost of temporal resolution, particularly when the interacting molecules are spatially stationary. Because the duration of observation directly relates to the NII, single-molecule observations for up to hours are possible, provided sufficient stage drift correction.

Notably, binding kinetics of individual antibodies have been previously resolved by using plasmonic nanoparticle biosensors. Localized surface plasmon resonance on functionalized nanorods has been utilized to reveal stochastic on/off rates that cause signal fluctuations after the equilibration of the interactions ^26^. SMIM confirms this observation as an abundance of transient, but specific interactions at equilibrium. Moreover, SMIM results indicate that such constant exchange of antibody molecules also take place in the cellular environment. As an added benefit, the incorporation of the PAINT principle enabled us to resolve the locations of antibody interactions with a localization accuracy on par with single-molecule localization microscopy.

While we have shown that the binding kinetics are heterogeneous, the local interaction dynamics are also governed by the local ligand concentration. In the case where primary β-tubulin antibodies represent the ligand, the local concentration is limited by the labeling density. STORM studies have revealed typical low labeling densities of antibodies, *i*.*e*., 10-30% ^30,31^, manifested as the discontinuous appearance of the microtubule fibers. The relatively low labeling density is attributed to the bulky size of the IgG antibodies and steric hindrance ^32^. In contrast, the 3xHA-α-tubulin cell line is likely to have a much higher local concentration of the HA tag. In addition to the higher expression of HA-tagged α-tubulin, the three HA tags on each α-tubulin increases the local ligand concentration and the on-rate of the 12CA5 similar to the repeating sequences and spacers employed by DNA-PAINT-ERS ^33^. This difference is consistent with the overall denser binding of 12CA5 compared to the Superclonal^™^ IgG at the same concentration (**Fig. 3a** *vs*. **2d**), aside from the different affinities between the antibodies.

The 12CA5 Fab displayed considerably lower on-rate compared to the 12CA5 IgG, indicating a significant reduction of the overall affinity of the 12CA5 Fab. The reduction of overall affinity is likely due to the monovalent interaction of a single paratope with an epitope in Fabs as opposed to bivalent interactions in IgGs. The removal of the Fc portion of IgG may also impact the stability and binding kinetics of the Fab molecule. Fc plays multiple roles in the formation and maintenance of the structure of IgG and also provides the flexibility to bind different sizes of antigens.

We obtained additional insights regarding the non-specific interactions using SMIM. Among the possible causes of non-specific interactions, one of the commonly accepted is the attraction of the Fc region of antibodies to endogenous Fc receptors on the surface of certain cells. Although it has been reported that FcRs do not retain the ability to bind the Fc portion after cell fixation ^34^, we observed non-specific binding events in all the experiments including the 12CA5 Fab without the Fc portion. SMIM maps of secondary to primary antibody binding illustrate that the use of normal goat serum as a blocking agent successfully reduced non-specific events from the secondary antibody raised in the goat. Similarly, using the mouse IgG2b kappa isotype control (the same isotype as 12CA5 IgG) in the blocking buffer reduced the non-specific binding of 12CA5-IgG to a greater extent (**Supplementary Fig. 9**).

Effective understanding of the antibody binding kinetics in the cellular environment provides a basis for the optimization of the therapeutic antibodies. At the single-molecule level, SMIM revealed highly heterogeneous and dynamic antibody-antigen interactions in the cellular environment. The observed binding kinetics were found to depend not only on the avidity and affinity, but also the local ligand concentration. SMIM paves the way for studying new classes of applications involving strong transient molecular interactions, such as ligand-receptor binding in the drug development and antibody neutralization capability in the vaccine development, one molecule at a time.

## Methods

### Materials

DMEM (11960069-500ml), penicillin-streptomycin (15140-122-100ml) L-Glutamine (25030-081-100ml) and 1x PBS (14190-144-500ml) were purchased from Gibco. FBS (F0926-500ml), puromycin (puromycin dihydrochloride; P8833-25MG), triton X-100 (9002-93-1-1L), MES (M3671-50G), EGTA (E3889-25G), sodium phosphate (342483-500G), DMF (227056-100ML), sodium bicarbonate buffer (S6014-500G) and cysteine-HCl (C1276) were ordered from Sigma. Normal goat serum (16210064, New Zealand origin), β-tubulin primary antibody (polyclonal, PA5-16863-500µl), goat anti-rabbit Alexa Fluor 647 IgG (Superclonal^™^ Recombinant, A27040-1mg), papain slurry (20341-5ml), Mouse IgG2b kappa Isotype Control (eBMG2b) eBioscience(tm) (14-4732-85), EDTA (AM9261-500mL), Protein A IgG binding buffer (21001) and Alexa Fluor(tm) 647 NHS Ester (Succinimidyl Ester; A20006) were purchased from Thermo Fisher Scientific. Potassium Chloride (7447-40-7-500G) and glycine (BP381-500) were obtained from Fisher Scientific. Ethanol (200 proof) was from Decon Labs Inc. Paraformaldehyde (16% w/v aq. stock solution; 50-00-0) was ordered from Alfa Aesar. Magnesium Chloride (7791-18-6-500G) was obtained from ACROS Organics. ReadyTag anti-HA (12CA5) (RT0268-25MG) was purchased from Bio X Cell. 8 well Chambered cover glasses (Borosilicate sterile, No 1.5, 155409) were ordered from Lab-Tek. Zeba(tm) spin desalting column (89889) and Pierce Protein A column (20356) were obtained from Thermo Scientific. Amicon® Ultra-4 Centrifugal Filter Units (30 K MWCO; UFC8030) were purchased from Millipore.

### Buffers

The following buffers were used for sample preparation and imaging.

- Cytoskeleton buffer: MES (10 mM, pH 6.1), Potassium Chloride (90 mM), Magnesium Chloride (3 mM) and EGTA (2 mM)
- Fixation buffer: Paraformaldehyde (3.7%), Triton-X-100 (0.5%), cytoskeleton buffer
- Blocking buffer for secondary staining: Normal goat serum (10%), Triton-X-100 (0.05%) in 1x PBS
- Blocking buffer for primary staining: BSA (5%) in 1x PBS, Mouse IgG2b kappa Isotype Control (10, 30, and 50 μg/mL) in BSA (5%)

### Cell culture

U2OS cells were cultured in DMEM supplemented with 10% FBS, 2mM L-Glutamine, and 100 units/mL penicillin-streptomycin. 3x-HA tubulin U2OS (clone 2E8 and 2F6) cells (referred to as mutant U2OS) were produced from a single cell clone and cultured in DMEM supplemented with 10% FBS, 2mM L-Glutamine, 100 units/mL penicillin-streptomycin, and 0.25 µg/mL puromycin. Both cell lines were maintained at 37 °C in a humidified atmosphere of 5% CO_2_ and split at confluence. For imaging, cells (∼10-20k) were seeded 1-2d before fixation in chambered coverglass slides.

### Sample preparation

For single-molecule imaging, samples were prepared in a chambered cover glass. Millipore water and ethanol cleaned chamber slides were air-dried and cleaned with plasma cleaner. Then, Superclonal^™^ goat anti-rabbit IgG-Alexa Fluor^™^ 647 (AF647) solution (10 pM) was incubated in the cover glass for ∼5 minutes. This solution was allowed to dry completely before the image acquisition. Once all the acquisition parameters were set, image acquisition started using a selected ROI. For SMIM imaging of secondary antibody staining, U2OS cells were seeded in a chambered coverglass and grown in an incubator under controlled conditions at 37 °C and 5% CO_2_. Following 24 hours of incubation, cells were washed twice with PBS and simultaneously fixed and permeabilized in the freshly prepared fixation buffer for 20 minutes at room temperature. Cells were washed thrice with PBS and were blocked for 30 minutes at room temperature on a rocker followed by incubation with primary antibody for β-tubulin (1 µg/mL in the blocking buffer) in a humid chamber for 2 hours at room temperature. After four rounds of washing steps with PBS, the secondary antibody staining solution containing Superclonal^™^ goat anti-rabbit IgG-Alexa Fluor^™^ 647 (AF647) (0.1 nM and 1 nM in blocking buffer) was added, and image acquisition was started. For SMIM imaging of primary staining, mutant U2OS cells were seeded, fixed, and permeabilized as with wild type U2OS cells, and were blocked for 2 hours with the corresponding blocking buffer at room temperature on a rocker. The primary antibody solution containing 1 nM 12CA5 IgG-AF647 in blocking buffer was added and image acquisition was started. For Fab staining, cells were prepared in the same procedure and 1 nM 12CA5 Fab-AF647 in blocking buffer was added for image acquisition. The same procedure was repeated with 0.1 nM concentrations of IgG-AF647 and Fab-647 solutions.

### Microscopy

Fluorescence imaging was carried out on an inverted microscope (Nikon Instruments, Eclipse Ti2). The single-molecule images were obtained with a laser power of 9.05 mW (measured after the objective) and an integration time of 50 ms. The 100x/1.49 objective was used with 1.5x external magnification. The Prime 95B cMOS camera was used at 16 bit without pixel binning. To optimize the NII duration, continuous acquisition, and different NIIs (0.5 s and 5 s) were used separately. For SMIM, 100 frames of images were collected with 5s NII using the same acquisition parameters.

### Data processing

A MATLAB program was developed based on the single-particle tracking program initially written by John C. Crocker^25^. A single-emitter algorithm was used to obtain the coordinates of each single molecule event. Single-molecule tracks were constructed using a maximum displacement distance of 220 nm between neighboring frames (pixel size = 110 nm), a minimum track length of two frames, and a maximum of two-time steps where the single molecule can be “lost” and “recovered” again. The coordinates from all events detected from each track were averaged to generate a single color-coded circle in the SMIM plots.

### Conjugation of Alexa Fluor NHS-ester to 12CA5 mouse IgG

ReadyTag anti-HA (12CA5) was adjusted to Ph 8.3 by adding 0.15 M sodium bicarbonate buffer to a final antibody concentration of 6.4 mg/mL. Alexa Fluor(tm) 647 NHS Ester was dissolved in DMF at 10 mg/mL. The antibody solution was added to the reactive dye solution at 20:1 dye to antibody molar ratio and was incubated for two hours at room temperature with continuous stirring. The dye was deactivated by adding 1 M Glycine (pH 7.4) to a final concentration of 30 mM. The reaction mixture was run twice through a Zeba(tm) Spin Desalting Column to purify the dye conjugated antibody from the excess dye. Antibody to dye labeling ratio was obtained using nanodrop measurements.

### 12CA5 Fab-Alexa Fluor 647 Production

#### Digestion of 12CA5 IgG-Alexa Fluor 647

The papain digestion of 12CA5-AF647 to generate Fab fragments is adapted from the manufacturer user guide at an enzyme to substrate ratio of 1:80. Well suspended aliquot of immobilized papain slurry was equilibrated in freshly prepared digestion buffer with 20mM Cysteine-HCl, 20mM sodium phosphate, and 10mM EDTA at pH 7.0 to activate the enzyme. The gel was separated from the buffer by centrifugation at 1000 g for 5 minutes and the washings were discarded. Equilibration was performed twice, and the gel was resuspended in a fresh volume of digestion buffer. The 12CA5-AF647 antibody was diluted to 0.625 µg/uL in digestion buffer and added into the resuspended immobilized papain slurry and incubated overnight (16 hours) in a shaker water bath at 37°C at high speed. The following day, 10mM Tris-HCl (pH 7.5) was added to the digest and the supernatant was separated by centrifugation at 1000 g for 15 min. Digestion was confirmed by performing an SDS PAGE.

#### Purification of 12CA5 Fab-Alexa Fluor 647

The supernatant from the previous step was run through a Pierce Protein A Column to purify Fab-AF647. All the steps were conducted in a dark room at room temperature. The protein A column was equilibrated with 5 mL of Protein A IgG binding buffer and allowed to drain by gravity. Supernatant from the previous step was diluted 1:1 Protein A IgG binding buffer and applied to the column and Protein A column was washed with another 15mL of the Protein A IgG binding buffer. Since the Fab fragments do not bind the column and are eluted with the flow-through, fractions of 0.5 µL were collected immediately after sample introduction. Absorbance at 280 nm was measured for each fraction with a nanodrop to monitor the protein elution with IgG binding buffer as the negative reference. Fractions with significant absorbances at 280 nm compared to IgG binding buffer were checked for the presence antibody by running an SDS-PAGE. Confirmed fractions were combined and concentrated using an Amicon® Ultra-4 Centrifugal Filter Unit.

### Nanodrop measurements

To measure dye-labeled protein samples, the “Protein and label” tab was selected from the home screen of NanoDrop One^c^ (Thermo Scientific) and specified the sample type and type of dye. 1-2 µL of blanking solution (PBS, 1X) was added to blank the instrument. Then, 1-2 µL of dye-labeled antibody solution was added onto the pedestal and started to obtain measurements. The Superclonal goat anti-rabbit IgG-AF647 sample was diluted in PBS (∼1:10) to obtain nanodrop measurements. The purified 12CA5-AF647 sample was diluted in PBS (∼ 1:10) to use for nanodrop measurements (**Supplementary Fig. 1 a,b)**.

## Acknowledgments

The authors thank the support from the Chicago Biomedical Consortium and the University of Illinois at Chicago.

## Statement of contributions

Y.S.H. conceived the project, H.G and T.P. performed the cell labeling and imaging experiments, H.G. performed antibody fragmentation and labeling, all performed data analysis and wrote the manuscript.

## Data availability statement

All data and MATLAB codes will be available upon request to the corresponding author.

