## Supplementary Figures 1-9 for "Single-molecule interaction microscopy reveals antibody binding kinetics"

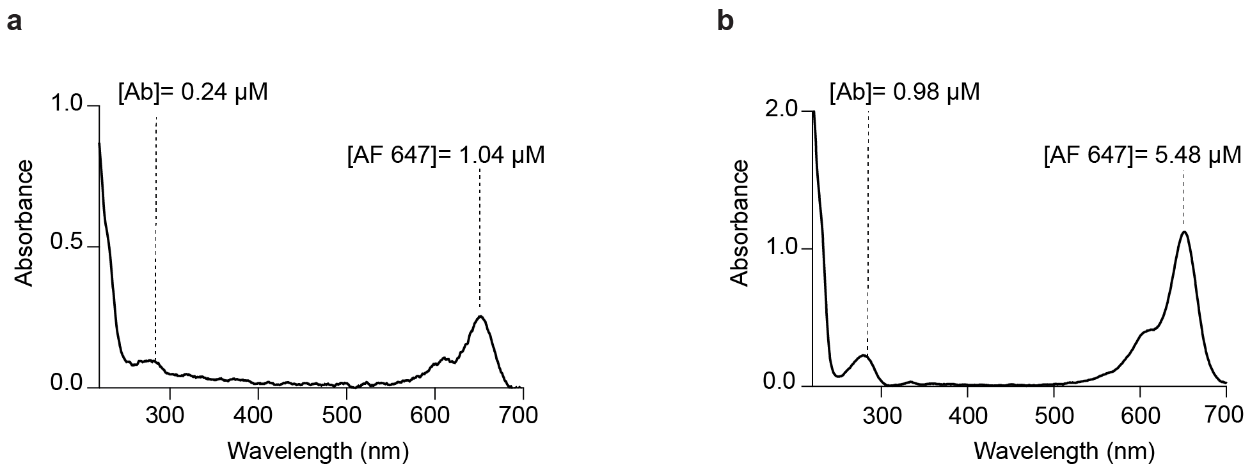


**Supplementary Fig. 1. Nanodrop measurements of dye-labeled antibodies.** The antibody: dye labeling ratio was calculated based on the measured antibody concentration [Ab] and dye concentration [AF 647]. **a**, Superclonal$^{TM}$ goat-anti-rabbit IgG-AF647 with the labeling ratio of 1:4. **b**, 12CA5 AF 647-IgG with the labeling ratio of 1:5.

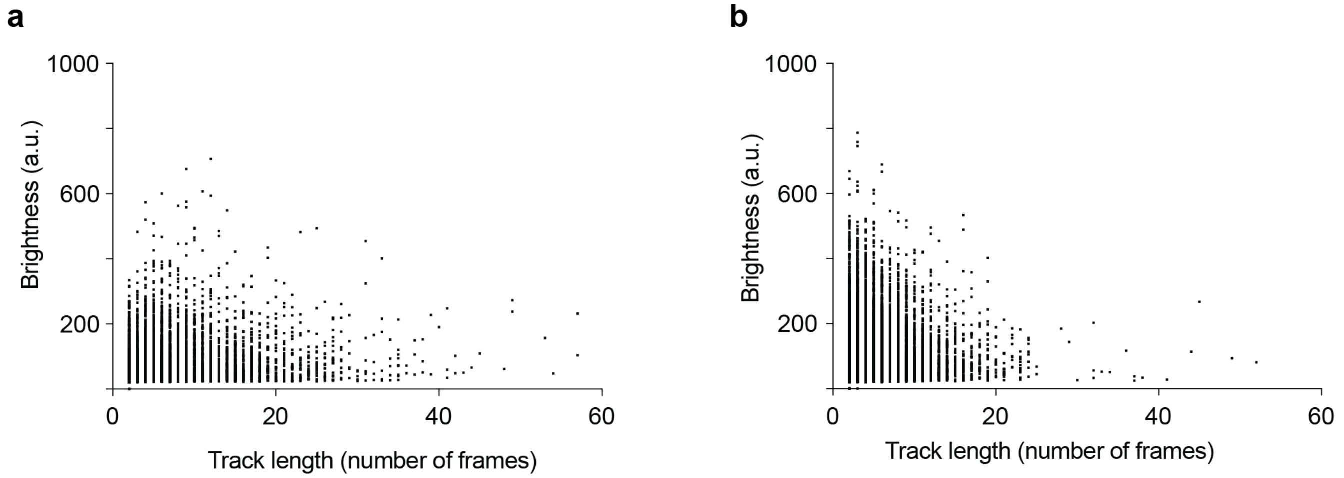


**Supplementary Fig. 2**. **Correlation between the event brightness and track length**. These plots show the correlation between the brightness and length of the tracks for the **a**, Superclonal$^{TM}$ goat anti-rabbit IgG-AF647 interaction with the primary antibody **b**, 12CA5 IgG-AF647 with interaction with HA. The dwell time is approximately 5 s multiplied by the number of frames.

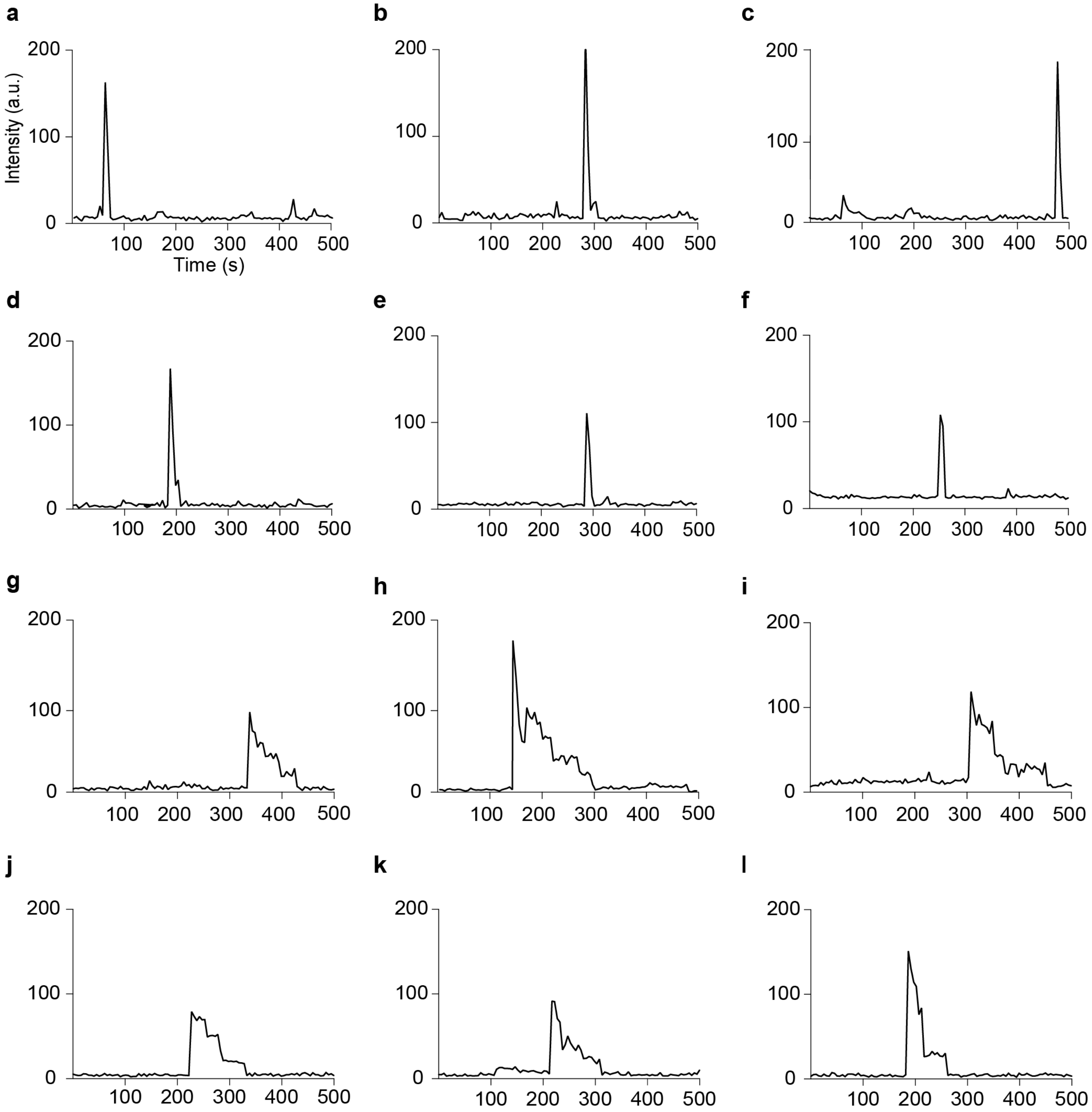


**Supplementary Fig. 3. SMAM revealed heterogeneous binding dynamics of Superclonal$^{TM}$ goat anti-rabbit IgG-AF647 antibody.** The representative intensity plots of transient (**a-f**) and stable (**g-l**) binding events of Superclonal$^{TM}$ goat-anti-rabbit IgG-AF647 in the presence of rabbit anti-tubulin primary antibody as the binding target.

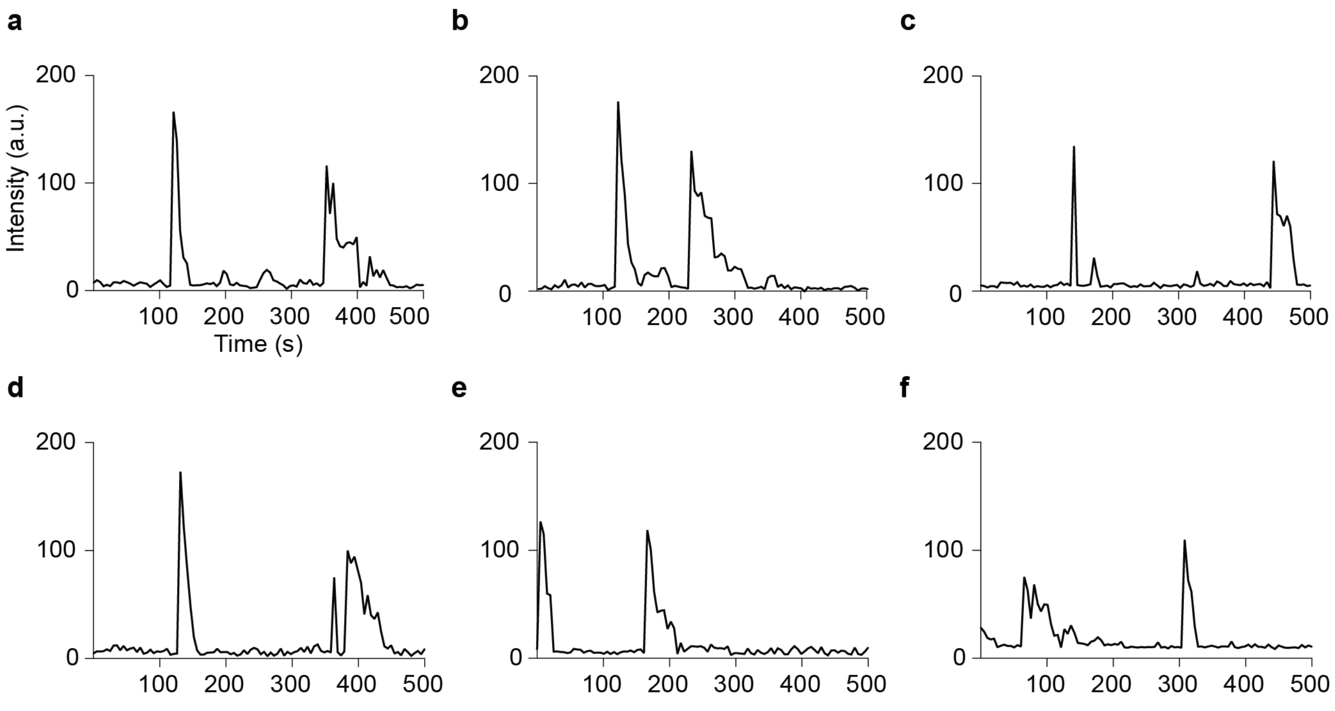


**Supplementary Fig. 4. Heterogeneous binding dynamics of Superclonal™ goat anti-rabbit IgG-AF647 antibody.** Multiple binding events within a diffracted-limited region of the interaction between Superclonal™ goat-anti-rabbit IgG-AF647 and rabbit anti-tubulin primary antibody.

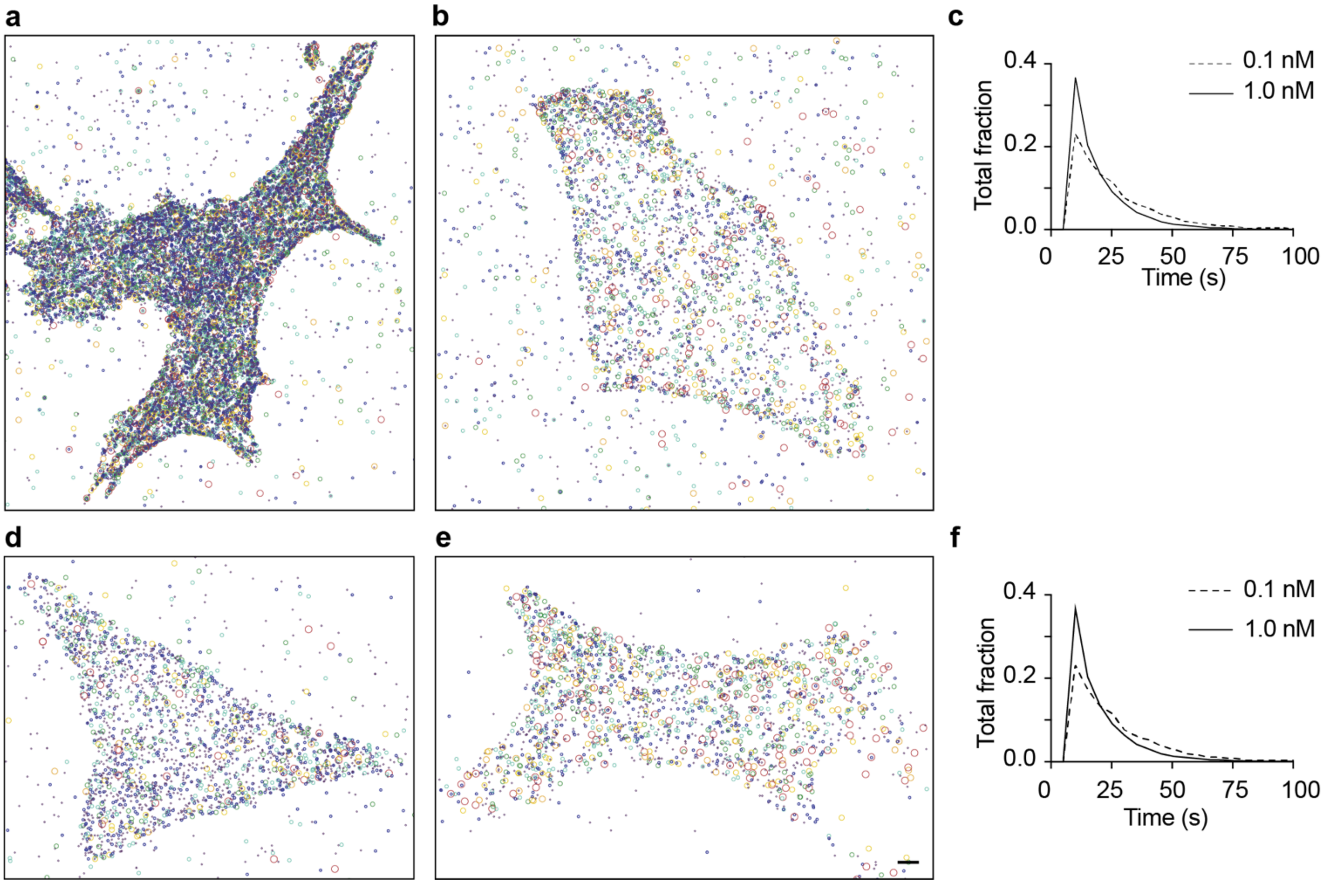


**Supplementary Fig. 5. SMIM interaction maps of 12CA5 IgG-AF647 and Fab-AF 647 reveal concentration-dependent kinetics**. SMIM interaction maps of primary staining with 12CA5 IgG-AF647 showing color-coded dwell times at, **a,** 1.0 nM and **b,** 0.1 nM in a fixed mutant U2OS cell (2F6) expressing 3xHA-α-tubulin blocked with mouse IgG2b kappa isotype in 5% BSA. **c,** Statistical comparison of binding events of **a** and **b**. SMIM interaction maps of primary staining with 12CA5 Fab-AF647 showing color-coded dwell times at, **d,** 1.0 nM and **e,** 0.1 nM in a fixed mutant U2OS cell (2F6) expressing 3xHA-α-tubulin blocked with mouse IgG2b kappa isotype in 5% BSA. **f,** Statistical comparison of binding events of **d** and **e**. Scale bar: 5 μm

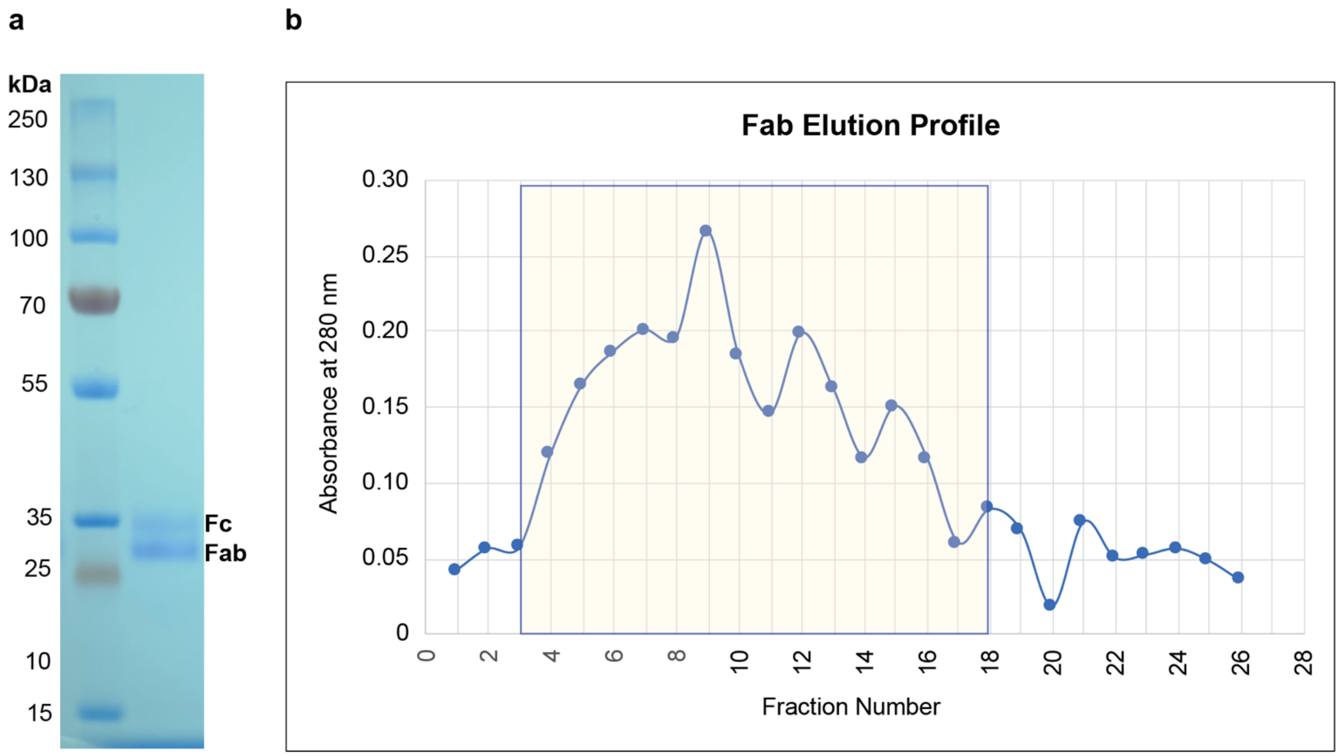


**Supplementary Fig. 6. 12CA5 Fab production using papain. a**, Papain digestion of 12CA5 confirmed on SDS PAGE showing bands corresponding to Fab and Fc fragments. **b**, Elution profile of Fab through Pierce Protein A Column to purify 12CA5-Fab. Fractions boxed in yellow were further confirmed for the presence of Fab by running an SDS-PAGE (not shown) and concentrated.

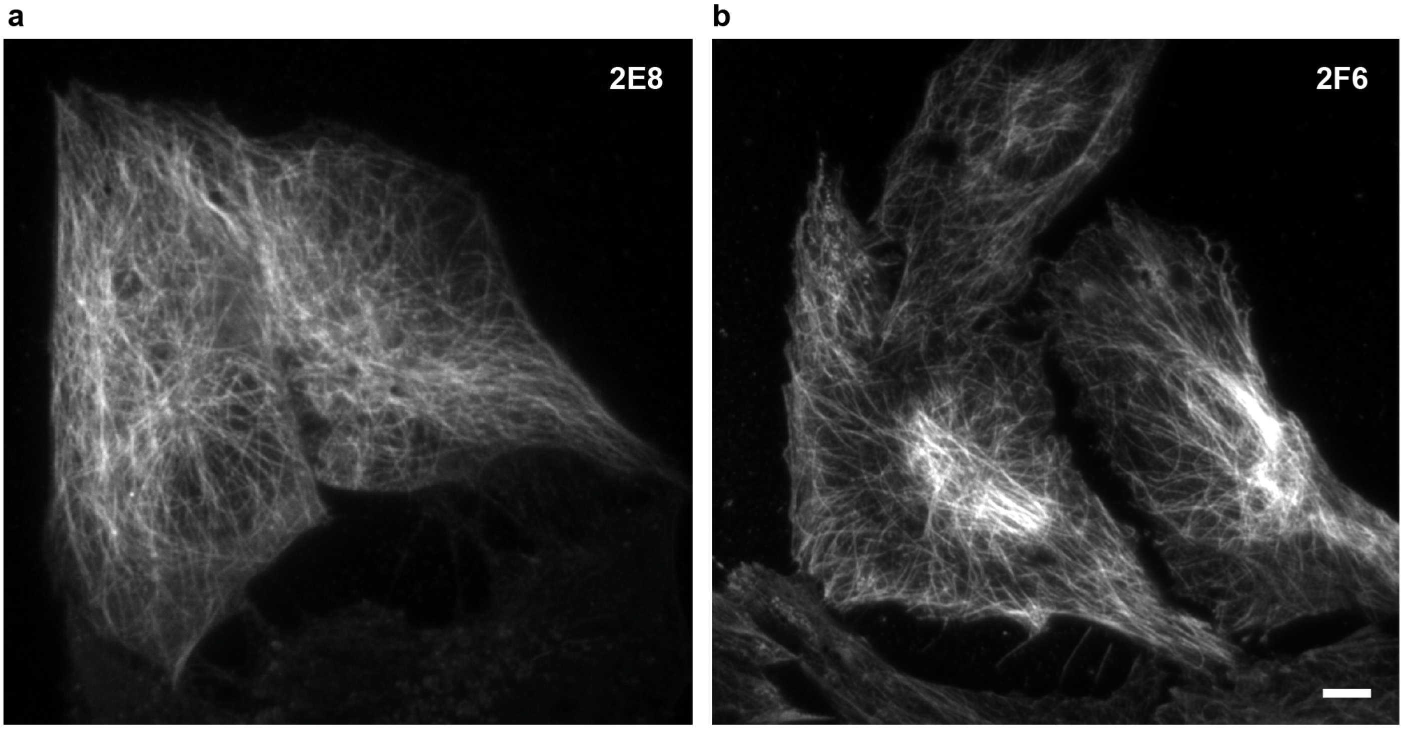


**Supplementary Fig. 7. Immunostaining using 12CA5-AF647 on the mutant U2OS cells expressing 3xHA on the N terminus of the α-tubulin**. **a**, 2E8 clone. **b**, 2F6 clone. Scale bar, 5 μm.

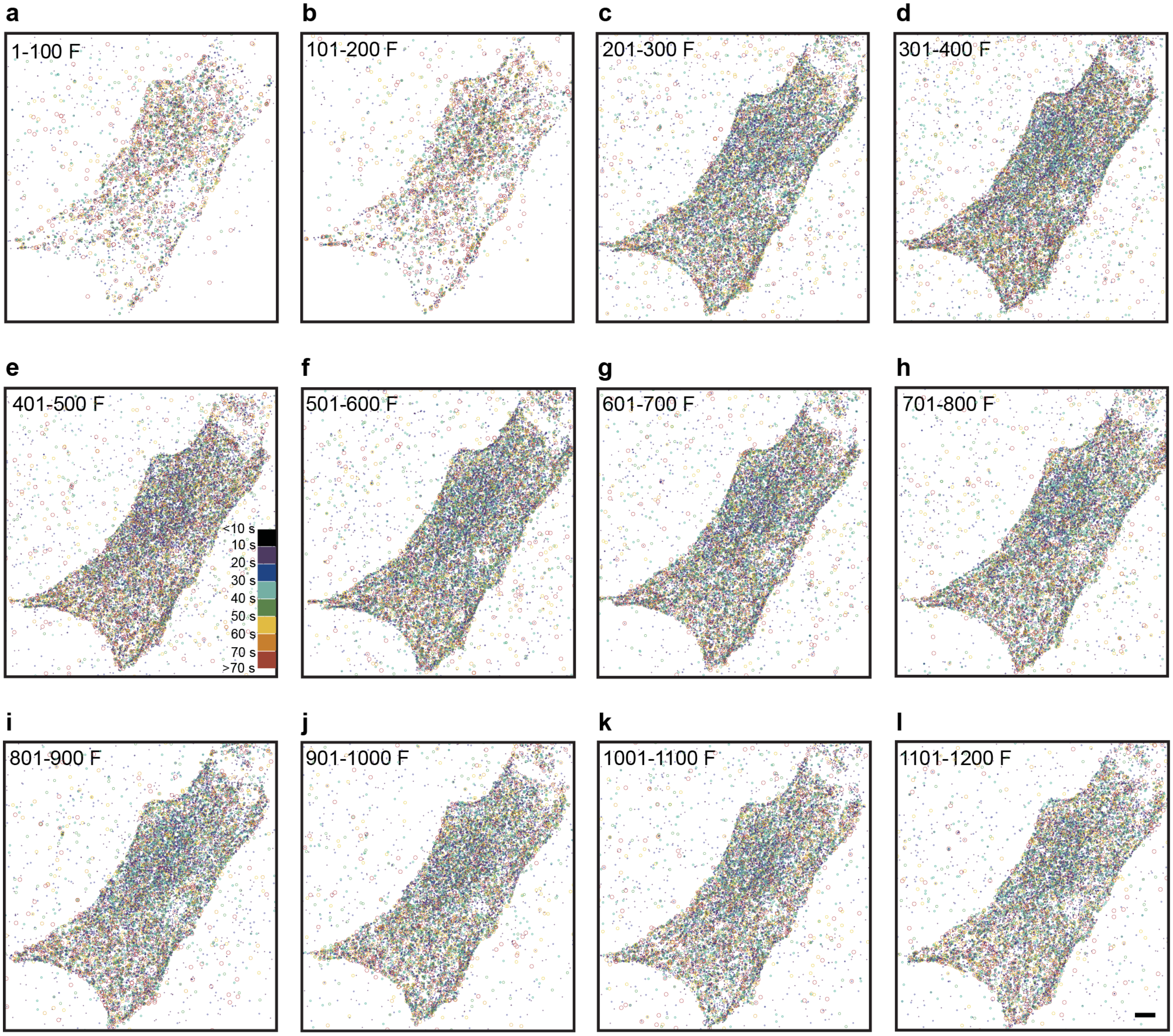


**Supplementary Fig. 8. A long-duration SMIM illustrates 12CA5 IgG-AF647 interactions with a fixed HA-expressing U2OS cell at 1.0 nM. a-l,** SMIM interaction maps of 100 frames each in a 1200-frame image acquisition of a fixed mutant U2OS cell (2E8) expressing 3xHA-α-tubulin, blocked with mouse IgG2b kappa isotype (10 ug/mL) in 5% BSA. Scale bar: 5 μm.

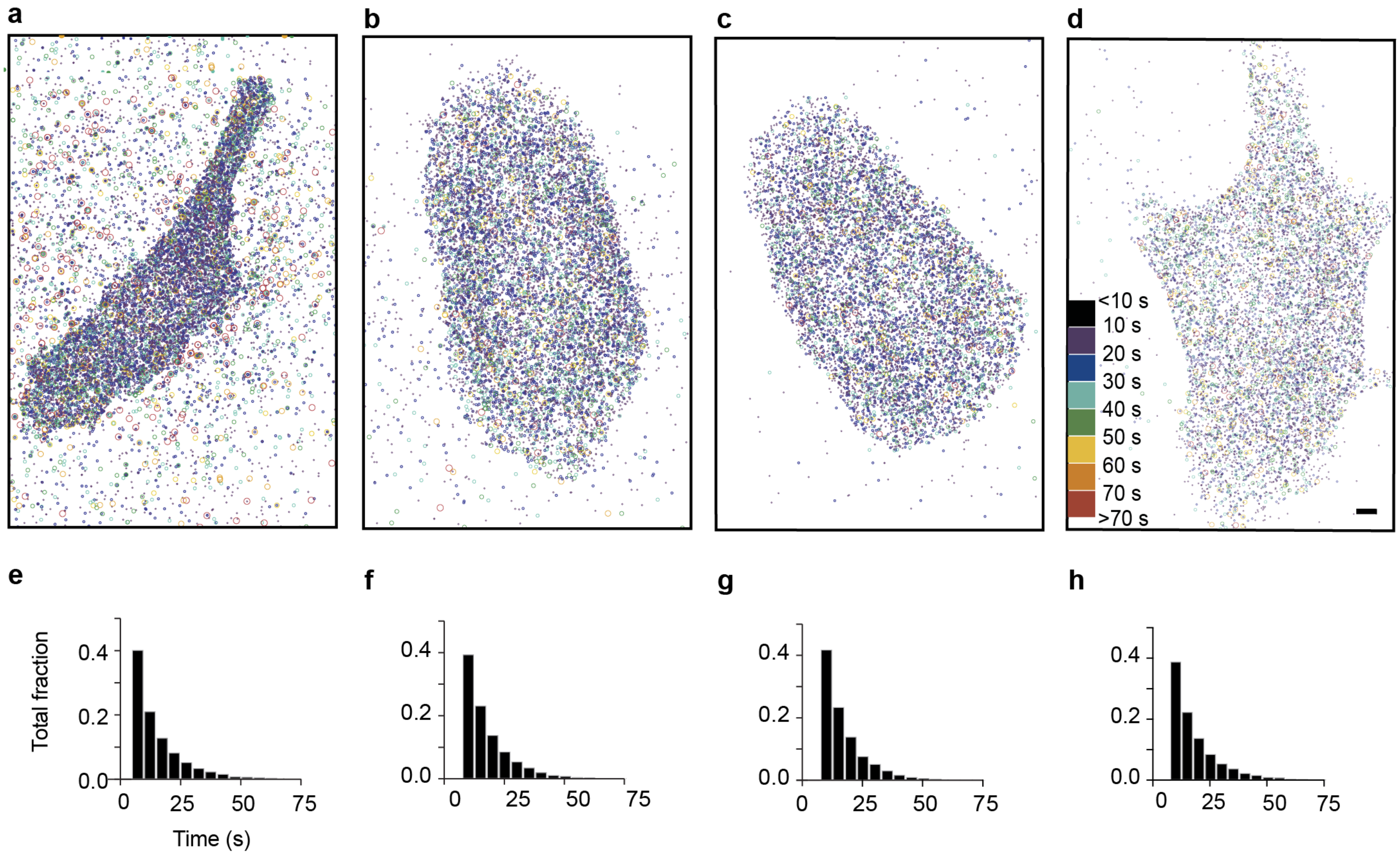


**Supplementary Fig. 9. SMIM on WT U2OS cells blocked with mouse IgG2b kappa isotype control shows the effects of blocking conditions on non-specific interactions of 12CA5 IgG/Fab-AF647.** SMIM interaction maps of 12CA5 IgG-AF647 at 1.0 nM on a wild type U2OS cell expressing 3xHA-α-tubulin blocked without the isotype, **a,** and with mouse IgG2b kappa isotype in 5% BSA at **b,** 10 µg/mL, **c**, 30 µg/mL, and **d**, 50µg/mL concentrations. **e-f,** Statistical analysis of binding events in **a-d** respectively. Scale bar: 5 µm
